# Influenza A Virus Infections Sense Host Membrane Tension to Dynamically Tune Assembly

**DOI:** 10.1101/2023.08.28.555166

**Authors:** Edward A. Partlow, Anna Jaeggi-Wong, Steven D. Planitzer, Nick Berg, Zhenyu Li, Tijana Ivanovic

## Abstract

Enveloped viruses often exhibit a pleomorphic morphology, ranging in size from 100nm spheres to tens-of-micron long filaments. For influenza A virus (IAV), spheres enable rapid replication and minimize metabolic cost, while filaments resist effects of antibodies or other cell-entry pressures. The current paradigm is that virion shape changes require genetic adaptation; however, a virus evolved to alter its shape phenotypically would outperform one that relies on genetic selection. Using a novel quantitative flow virometry assay to characterize virion shape dynamics we find that IAV rapidly tunes its shape distribution to favor spheres under optimal, and filaments under attenuating conditions including the presence of antibodies. We identify membrane tension as a key cue sensed by IAV determining shape distributions. This phenotypic shift outpaces genetic change and serves to enable additional life cycles under pressure. Our work expands knowledge of the complex host-virus interplay to include viral responses to the local environment by optimizing its structure to maximize replication and ultimately host-host transmission.

## Introduction

Influenza A virus (IAV) is a major public health concern, with both regular seasonal outbreaks and occasional pandemics. IAV is a member of the *Orthomyxoviridae* family, characterized by a segmented negative-sense RNA genome and the major surface glycoproteins hemagglutinin (HA) and neuraminidase (NA). Influenza is pleomorphic; virions can adopt a range of morphologies, from small 100 nm spheres to filaments with lengths of several microns. Pleomorphy is not unique to influenza, but is employed by many enveloped virus families, despite extreme diversity in structure, composition, and entry strategy. Many of these pleomorphic viruses, such as respiratory syncytial virus^1^, measles virus^2^, Ebola virus^3^, Nipah virus, and Hendra virus^4^ pose great threats to human health.

The emergence of filamentous morphologies across viral phylogenies, despite a greater demand for cellular resources to build larger particles, implies that viruses may generally benefit from this type of assembly. Indeed, for influenza, filaments are selected for in animal infections but not in tissue culture^5–8^. However, the mechanisms by which influenza adopts its morphology are not clear. Attempts to explore the biology of filaments have been limited by multiple challenges. Measurement of particle shape by electron microscopy is quantitative but requires extensive sample concentration and processing, and the throughput is limited to hundreds or thousands of particles. There is also risk that the processing steps enrich certain virion morphologies. Additionally, large filaments can be fragile, and processing steps can lead to fragmentation and loss of these particles. Nevertheless, multiple mutations that influence influenza shape have been characterized^9–16^.

Much focus on the assembly of influenza filaments has been on the matrix protein, M1, which coats the lumen of virus particles. The M1 gene segment of A/Udorn/72 is sufficient to confer a filamentous phenotype to other strains^9^. Additionally, mutations in the M1 segment have been found to alter the morphology of assembled virions^9–12^. Electron microscopy of influenza shows that M1 assembles into an ordered, helical assembly in filaments but not in spheres where M1 appears disordered^13,14^. These observations have led to a model where M1 oligomerization competes with viral fission. M1 variants that more readily oligomerize would therefore be expected to produce a more filamentous population.

Mutations affecting shape have also been reported in NA^15^ and nucleoprotein (NP)^16^. Electron microscopy and superresolution microscopy have been applied to study the organization of these viral components in filaments. In some filaments, the ribonucleoprotein complex (RNP) containing the genome is packaged at one end of the filament^13^, however this is not universal to all filaments^17^. Some filaments terminate in a large bulbous region called an Archetti body^18^, which houses the RNP and is enriched in the NA glycoprotein^17^, while others have no apparent organization of RNP or either glycoprotein. Overall, the mechanisms of how NA and NP mutations alter shape remain unexplained.

Other work has attempted to explain the pressures that select for filaments in animal infections^7,8,10,19^. Several models propose that the organization of glycoproteins in filaments could confer advantages in the context of an animal infection. NA enrichment at one end of a filament can promote directional diffusion through a sialic acid-rich mucous^20^. Additionally, it has been proposed that the ratio of NA to HA varies with filament length, and that certain subsets of particles could stand to tolerate inhibition of these proteins, which must work in a balance for successful infection^21^. Our previous work has shown that filamentous influenza virions and Ebola virus-like particles have a direct advantage to attachment and fusion under antibody challenge^22^. A larger contact area increases the likelihood of successful attachment and subsequent fusion when glycoprotein activity is compromised, such as by antibody neutralization, introduction to a new host, or by spontaneous attenuating mutations^22^. However, mutations that increase filaments might be less efficient than mutations that directly bypass the challenge, such as an antibody-escape mutant. It thus remains unexplained why filamentous influenza is ubiquitous in animal infections.

Factors that select for filaments, such as presence of antibodies or introduction to a new host, can arise on time scales shorter than the viral replication cycle. Current models suggest that genetic selection drives the emergence of more filamentous variants during *in vivo* infections, however the coincidence of a filament-producing mutant with such a challenge may be improbable, particularly during the genetic bottlenecks typical of viral propagation within and between hosts^23^. A strain might need to survive multiple generations under these pressures before a mutation promoting filament production arises. In contrast, a phenotypic shift to a filamentous morphology in response to external pressures would be immediately beneficial to infection, but the possibility that shape can be dynamically tuned is unexplored.

Here, we show that influenza morphology is dynamic during infection. Using high throughput flow virometry performed directly on minimally processed infection supernatants, we find that influenza infections rapidly change their shape distribution in response to factors typical of animal infections, such as low input virus or the presence of antiviral antibodies. We further show that virion assembly favors filaments in response to treatments that increase host membrane tension, suggesting a mechanism by which influenza monitors and responds to the environment during infection. This phenomenon of real-time adaptive changes to an infection in response to attenuating pressure adds a new dimension to our understanding of viral immune evasion and persistence strategies.

## Results

### Influenza A shape distribution is dynamic

Determining whether viral shape changes in response to environmental challenges requires the ability to measure low concentrations of virions produced under inhibitory conditions. Here we use ‘virions’ to refer to any viral particle (based on presence of surface HA) regardless of whether it is solely competent for infection. To determine both overall virion concentration and the relative proportion of filaments, we employed flow virometry to measure the violet side scatter (VSSC) of virions (Figure 1A). We used samples diluted from high concentration of purified virions to enable their correlation to electron microscopy (EM), which requires high sample concentrations. Nearly all virions produced detectable scatter over background, and, moreover, filamentous particles scattered violet light more strongly than the spherical ones (Figure 1A). Consistent particle counts were obtained by flow virometry, both with and without normalization to reference beads, and were correlated to hemagglutination assays with a scaling factor of 8.9 x 10^5^ particles/HA unit (Figure S1D). This is in line with previously reported values^24–26^. Flow virometry also revealed two distinct subpopulations in virion samples that were enriched for filaments (Figure 1B, S1B). The population with greater VSSC from flow virometry correlated to filamentous particles longer than 130nm, as observed by negative-stain electron microscopy (Figure 1B, Figure S1A-C). In sum, established assays - HA assay for particle counts, and EM for particle shape - validated this novel approach for obtaining fast yield and shape measurements of influenza A virions.

**Figure 1.**
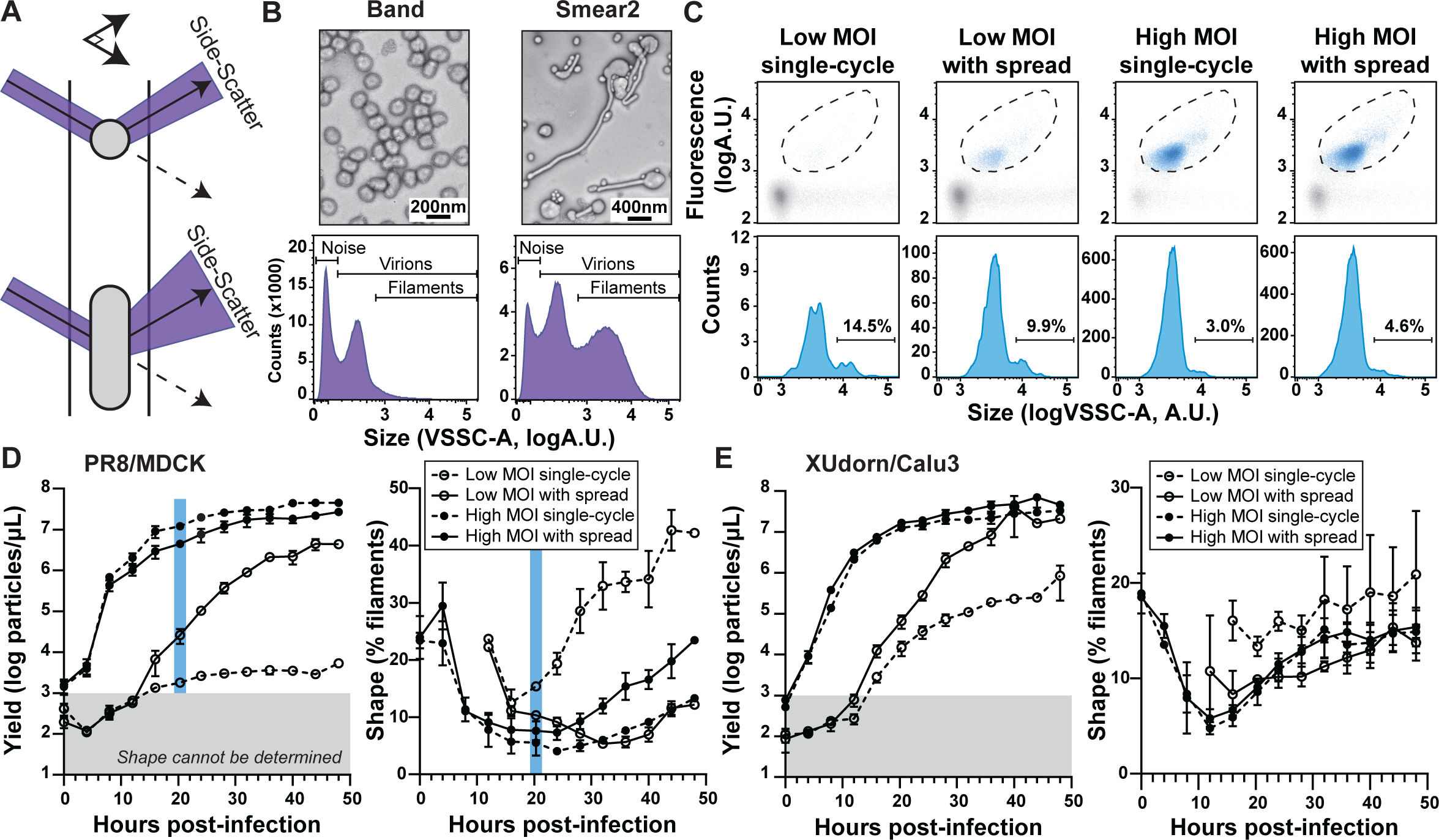
Influenza A shape distribution is dynamic. **A**. Left: A schematic illustrating the violet side-scatter (VSSC) of particles of varying sizes. Large particles such as filaments scatter violet light more than smaller particles such as spheres. **B.** Top: Electron micrographs of samples enriched in spherical (Band) and filamentous (Smear2) Influenza A virions by sucrose gradient ultracentrifugation. Bottom: Corresponding VSSC histograms of above samples. Violet side-scatter of virions is mostly resolved from noise inherent to small-particle flow cytometry. **C**. Representative flow virometry data for unpurified infection supernatants. To distinguish highly dilute virus signal from noise, virions are treated with a Dylight 550-labeled HA antibody (Sb H36-26) to enable separation in two detection channels. Displayed are scatterplots of VSSC versus fluorescence with gates enclosing viral particles (dashed line, top), along with corresponding VSSC histograms of the gated population (bottom). Percentage of filamentous virions for each sample are indicated. **D**. Virion yields (left) and shape (right), determined by flow virometry of supernatants from MDCK cells infected with PR8 at MOI 0.006 (Low MOI) or MOI 6 (High MOI). Single-cycle infections were achieved by omitting trypsin. Mean and S.E.M. are plotted. The blue bars highlight the samples with raw data shown in panel C. Gray shading indicates yields too low to reliably measure shape. **E**. Similar to D, except for Udorn infections in Calu3 cells.

To extend the sensitivity of flow virometry measurements to dilute samples, including low-yield culture supernatants, we used a fluorescently labeled antibody targeting the virion surface protein, HA. This treatment helped separate virions from the inherent noise present in small-particle flow cytometry by enabling gating in both fluorescence and VSSC channels (Figure 1C). This labeling also revealed stronger HA intensity for viral particles with greater VSSC, as expected for larger virions. As a result, we were able to achieve count and shape measurements for samples as dilute as 1000 particles/µL. This concentration is typical for a majorly attenuated infection (corresponding to ∼1HA unit/ml, Figure S1D), and is approximately 100,000 times lower than the concentrations required for electron microscopy (Figure 1B, S1A)^22^.

We used flow virometry to measure virus yield and shape over the time course of infections using two influenza A strains, A/Puerto Rico/8/34 (PR8) and a variant of A/Udorn/72 (Udorn) containing the HA segment of A/Aichi/2/1968-X31, herein called XUdorn. PR8 is reported to be spherical, while Udorn is reported to be filamentous^27^. Replacing the HA segment of Udorn with that of A/Aichi/2/1968-X31 does not affect the shape properties of Udorn, which are determined by its M1 segment. We performed the time course analysis in two cell lines, MDCK and Calu3. We infected cells at two multiplicities (MOIs), MOI 6 and 0.006 infectious units (IU) per cell. These inputs were chosen to create conditions where either 1) nearly every cell is infected by multiple infectious units (High MOI), or 2) each infected cell receives only one infectious unit (Low MOI) (Figure S2). To limit infections to the initially infected cells, we omitted trypsin protease, which is required for the activation of HA on produced virions to enable further rounds of infection^22^. We also inhibited spread by the addition of ammonium chloride 4 hours post infection, which alkalinizes organelles and blocks pH-mediated membrane fusion of influenza in the endosome (Figure S4). These controls confirmed that omitting trypsin effectively inhibits spread in MDCK cells, ensuring single-cycle infections (Figure S4), although we saw that Calu3 cells permit some spread even in the absence of trypsin, in line with previous reports^28^.

Without trypsin, the rate of virion production remained relatively constant until late in infection, and was lower for the low MOI cases for all time points in infection (Figure 1, S5). In contrast, when trypsin was included in low MOI samples, the rate of virion production increased, and the virion yields approached the High MOI values, over the course of infection due to produced virions infecting new cells. Notably, we found that the shape distribution of the produced virions was dynamic over the course of infection. Infections generally produced a somewhat filamentous population first, followed by a period dominated by spherical particle production, with an eventual increase in filament production as time progressed. With PR8 in MDCK cells, we also found that low MOI single-cycle infections produced a more filamentous population than high MOI or low MOI infections with spread (Figure 1). We also reproduced previous findings that XUdorn produces a high proportion of filaments in MDCK cells, and observed this at both MOIs tested with and without spread (Figure S3B). In contrast, XUdorn was no more filamentous than PR8 in Calu3 cells (Figure S4), an unexpected finding that highlights the extent to which the cellular environment of infection can direct viral assembly. In summary, we have developed and validated a flow virometry assay for high-sensitivity virion count and shape measurements including for majorly attenuated infection. We used this novel assay to identify the stage of infection, MOI, and cell line as non-genetic determinants of virion shape. These combined data show that the shape distribution of assembling virions is not genetically fixed as previously assumed, but is dynamic over infection.

### Infections with lower virion yield produce more filaments

To probe the effect of MOI on shape, we sought to determine the relationship between input virus amount and burst size, or the average number of virions produced per infected cell. We first measured burst from the single-cycle infection data in Figure 1 (Figure 2A). Burst sizes were larger at high MOI than low MOI correlating to virion production rates at these MOIs (Figure S5), though the relative ratio of burst sizes varied depending on the cell type. Specifically, we observed ∼15-fold increase in PR8 particles (from 1700 to 25,000) and ∼7-fold increase in XUdorn particles (from 1600 to 11,000) produced per infected MDCK cell at high MOI compared to low MOI. In Calu3 cells, the ratios of burst sizes for high to low MOI were lower at ∼2.0-fold for PR8 and ∼2.5-fold for Calu3. This mostly was an outcome of greater yields in Calu3 cells at the low MOI. Notably, MOI only significantly affected shape when there was the largest difference in burst size (PR8 in MDCK), implicating some relationship between viral production and shape (Figure 2). An exception was XUdorn infection of MDCK cells where all conditions yielded a high proportion of filaments and where a switch to greater filament production occurred much earlier over the infection time-course (Figures S3 and S4). In this case, cell-type effects appeared dominant over effects from input and burst size.

**Figure 2.**
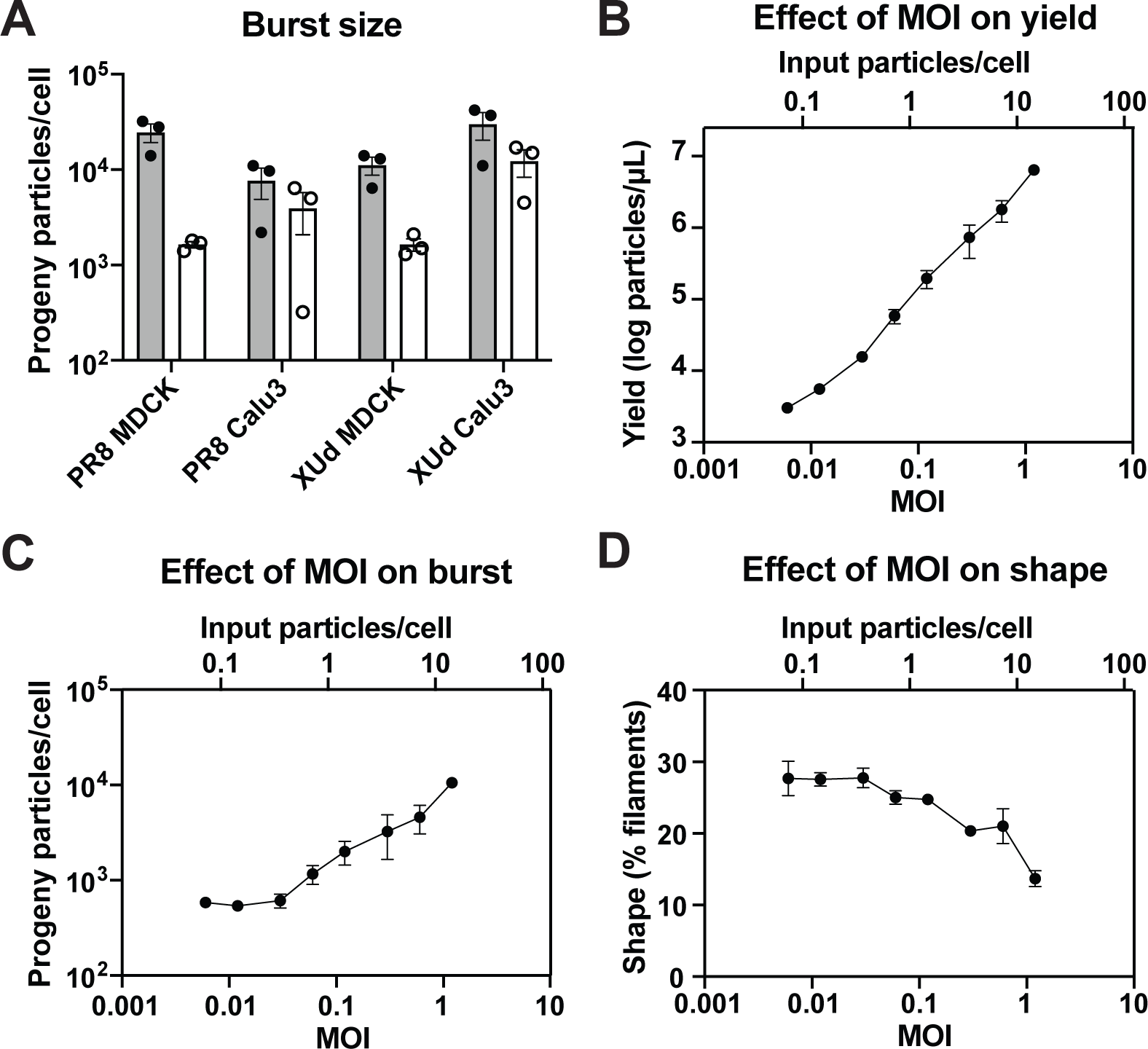
Burst Size Influences Virion Shape. **A.** Burst sizes (virions produced per infected cell) were determined from the time courses in figure 1 by dividing cumulative yields at the final time points by the expected number of infected cells from Poisson distribution (see Figure S2). **B**. Yields measured at 24 hours post infection by flow virometry of supernatants of MDCK cells infected with PR8 at various MOIs. **C**. Burst sizes of the samples in B calculated by dividing total yield displayed in panel B by the expected number of infected cells from Poisson distribution. **D**. Shape of the samples in B measured by flow virometry. All plots show mean and S.E.M.

To explore burst size effects further, we infected MDCK cells with PR8 without trypsin at various MOIs and quantified virion production and shape at 24 hours post infection (h.p.i.). As expected, virions produced increased steadily as the MOI increased (Figure 2B). We found that burst size and virion shape, however, were constant at very low MOI (≤0.06), where any infected cells are expected to have received only one infectious unit, and ∼16% of cells received multiple particles, based on a particle/pfu ratio of 12 as determined by flow cytometry (Figure 2C, D, S2A, B). As MOI increases further, the extent of infection for these particle inputs might thus be greater than predicted from the infectivity data alone. An infection may be enhanced through gene segments contributed by a noninfectious virion, or more cells can become infected due to complementation from coincident entry by incomplete virions^29,30^; thus the ratio of output to input virion amounts increases nonlinearly at inputs where coincidence of particles is expected. Increase in the MOI resulted in a corresponding increase in burst size and decrease in production of filaments. Thus the viral load within a cell determines the morphology of its produced virions. Together, the experiments in Figures 1 and 2 indicate a direct correlation between virion output and virion shape, with a higher production reducing, and lower production promoting, filamentous virion assembly.

### Antibody pressure drives filament assembly

Our previous work found that filaments have a replicative advantage when neutralizing antibodies are present^22^. We therefore tested whether influenza shape varied in response to antibody pressure. We infected MDCK cells with PR8 in the presence of varying doses of a panel of antibodies specific for viral surface antigens and measured virion yield and shape after 24 hours. The panel included antibodies targeting seven distinct antigenic regions across all IAV surface proteins: four in the HA head domain and one each in the HA base, NA, and M2. Strikingly, all 12 antibodies elicited a significant increase in the production of filaments, regardless of whether the antibody effectively neutralized infection (Figures 3, S6). We also included samples where antibody was present only during viral attachment to cells and the first 4 hours of infection (PRE), and samples where antibody was only present after 4 hours when viral internalization is expected to be complete (POST). We found that pre-treatment elicited shape effects when infection was attenuated, but post-treatment effects were generally stronger and independent of neutralization. Importantly, these antibody-induced filaments retain their infectivity, exhibiting similar per-particle infectivity as the input virus or the supernatants of untreated controls in the same experiment (Figure S9). This further underscores the impact of antibodies on virion shape without compromising their infectious potential.

**Figure 3.**
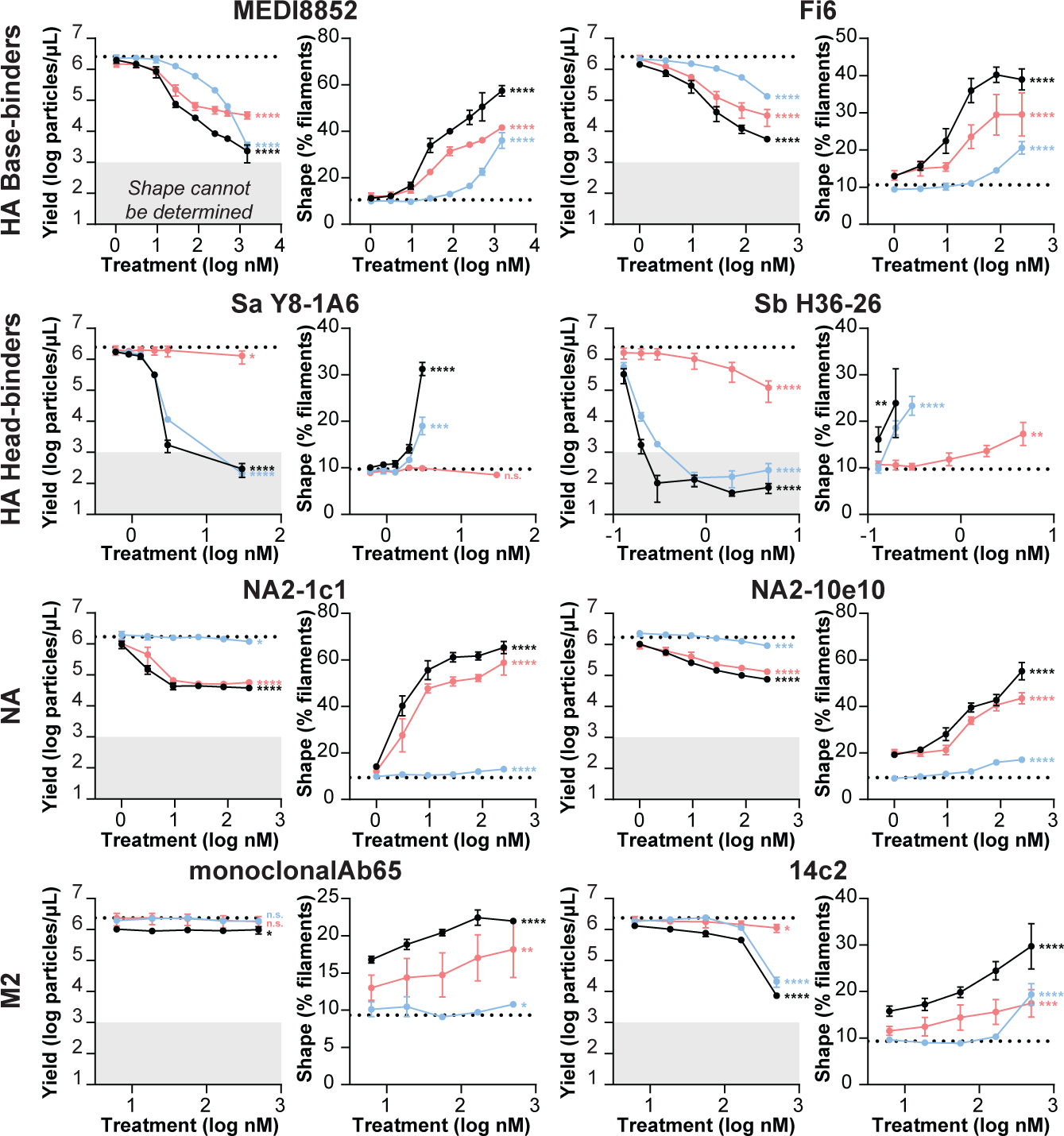
Antibody Pressure Drives Filament Assembly. Plots of viral yields (left) and shapes (right) measured by flow virometry of supernatants from MDCK cells, 24 hours post-PR8 infection at MOI 0.6, treated with antibodies. Black: antibody present throughout infection. Blue: antibody present until 4 hours post-infection to test entry effects. Red: antibody added at 4 hours post-infection to test assembly effects. All antibodies are mouse IgG except Fi6, which is human IgG. All plots show mean and S.E.M. *p<0.05, **p<0.01, ***p<0.001, ****p<0.0001 by unpaired t-test relative to an untreated control for the highest antibody concentration. Gray shading indicates yields too low to reliably measure shape.

We repeated this experiment with PR8 in Calu3 cells (Figure S7) and with XUdorn in Calu3 cells (Figure S8). For XUdorn, a different but overlapping set of 9 antibodies was used because of strain antigenic differences. This group included TriHSB.2, a computationally designed influenza neutralizing protein that targets the HA receptor-binding domain^31^. In Calu3 cells, continuous-treatment (ALL) and post-treatment elicited shape effects, although weaker than those seen in MDCK cells. Additionally, there were minimal effects of pre-treatment on virion shape in Calu3 cells, in line with how change of MOI did not affect shape in this cell line. Overall, shape is dynamically tunable in the presence of external pressures, both by reducing the viral replication (lower MOI or antibody pre-treatment) or by another mechanism during assembly (antibody post-treatment). This effect is general to antibodies of various targets, although the magnitude of changes grouped by antigenic region. Finally, the response is conserved in multiple IAV strains in multiple cell lines, though cell/strain differences in extent are apparent.

### Shape dynamics are fast and specific

To determine the kinetics of antibody-induced shape changes, we performed a short time-course of shape measurements after addition of antibody to an already established infection. We chose to start the antibody treatment 20 hours post-infection of MDCK cells with PR8 virus because this is when shape is most stable in our time courses (see Figure 1D). At the time of treatment, cells were washed to remove any already produced virus, and given either fresh media, or media with various concentrations of MEDI8852 antibody. As expected, virion shape was fairly constant in untreated media, slowly increasing from 7.5% to 10.1% over the 4-hour period while yield steadily increased (Dotted line, Figure 4A, B). In contrast, antibody treatment elicited a rapid and dose-dependent increase in the population of filaments without significantly changing viral yields (Figure 4A, B). These data further support that antibodies directly affect the assembly of viral particles independent of their ability to neutralize infection. Additionally, the rapid nature of these dynamics likely excludes genetic or gene expression changes as the underlying mechanism.

**Figure 4.**
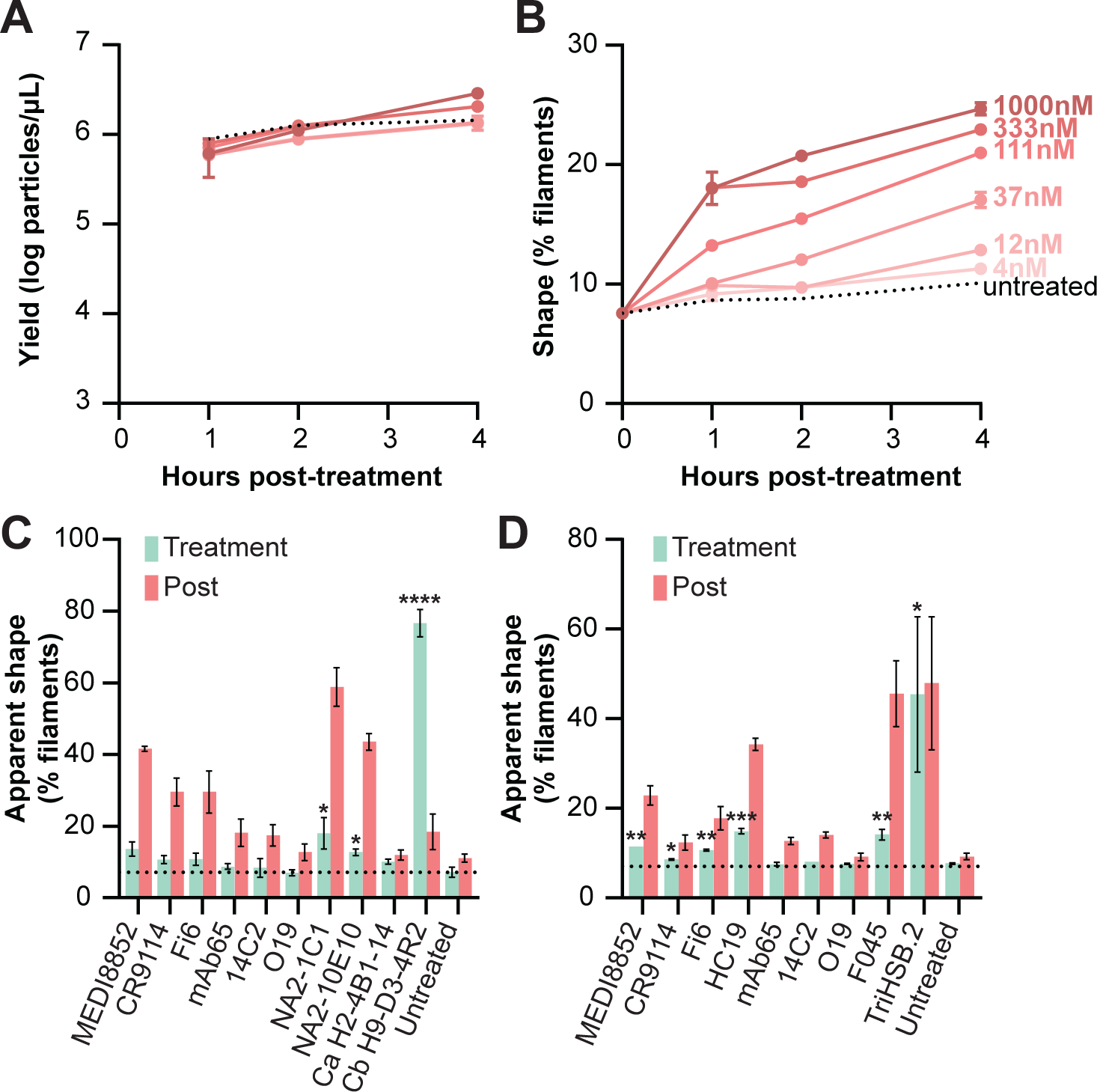
Shape Dynamics are Fast and Specific. MDCK cells infected with PR8 at MOI 0.6 for 20 hours were washed into media containing varying concentrations of MEDI8852 antibody. **A**. Viral yields during 1, 2, or 4 hours following antibody treatment. **B**. Shape of the samples in A. **C**. Compares apparent shape of supernatants from untreated infections mixed with antibodies (green) to the shapes produced in infections treated with the same antibody during assembly (red) in Figures 3 and S5. mAb65: monoclonalAb65. **D**. As in C, but for Udorn relative to the post-treatments in Figure S7. All plots show mean and S.E.M. *p<0.05, **p<0.01, ***p<0.001, ****p<0.0001 by unpaired t-test relative to an untreated control.

Some antibodies induce clumping of virions by simultaneously binding to multiple virions^32^. Clumped virions can appear similar to filaments by flow virometry in both fluorescent and VSSC channels. For all antibodies used, we tested whether their addition to supernatants from untreated infections affected virions’ apparent shape by flow virometry. While some statistically significant changes to shape distributions were observed, they were too minor to explain the changes observed in treatment experiments, except for Cb H9-D4-4R2 antibody and TriHSB.2 trimeric binder (Figure 4C, D). These results, along with observations that particle numbers need not decrease to observe shape changes (Figure 3), support that our findings are due to differences in newly assembled particle shape rather than antibody-induced clumping.

### Filament assembly responds to membrane tension

The rapid nature of antibody-induced changes in viral assembly could implicate a biophysical effect driving filament assembly. To that end, we aimed to test how virion assembly responds to general changes to membrane biophysics. We subjected MDCK cells infected with PR8 to osmotic shock at 20 h.p.i. and measured the shape of viral particles produced in the following 30 minutes. Hypotonic media induces cell swelling by osmosis, increasing membrane tension, while hypertonic media has the opposite effect. Hypotonic media generated populations greater than 50% filaments while hypertonic media suppressed filaments to below 10% (Figure 5B). Interestingly, we also saw small but significant changes in the number of particles produced, increasing ∼2-fold in hypertonic media and decreasing ∼2-fold in hypotonic (Figure 5A). We reinfected MDCK cells with equivalent numbers of each of these particles (3 particles/cell) and found little difference in per-particle infectivity or the shape of virions produced from these infections (Figure 5C, D). In summary, osmotic changes alter virion assembly and output. Despite their altered morphology, these particles are authentic virions, as the treatment does not influence per-particle infectivity or virion shape in subsequent infections. Additionally, the rapidity of these changes and the failure of phenotypes to persist to future generations further supports a mechanism for shape adaptation that does not require genetic change. However, from these results alone, we cannot distinguish whether membrane tension or media salinity drive these changes in virion assembly.

**Figure 5.**
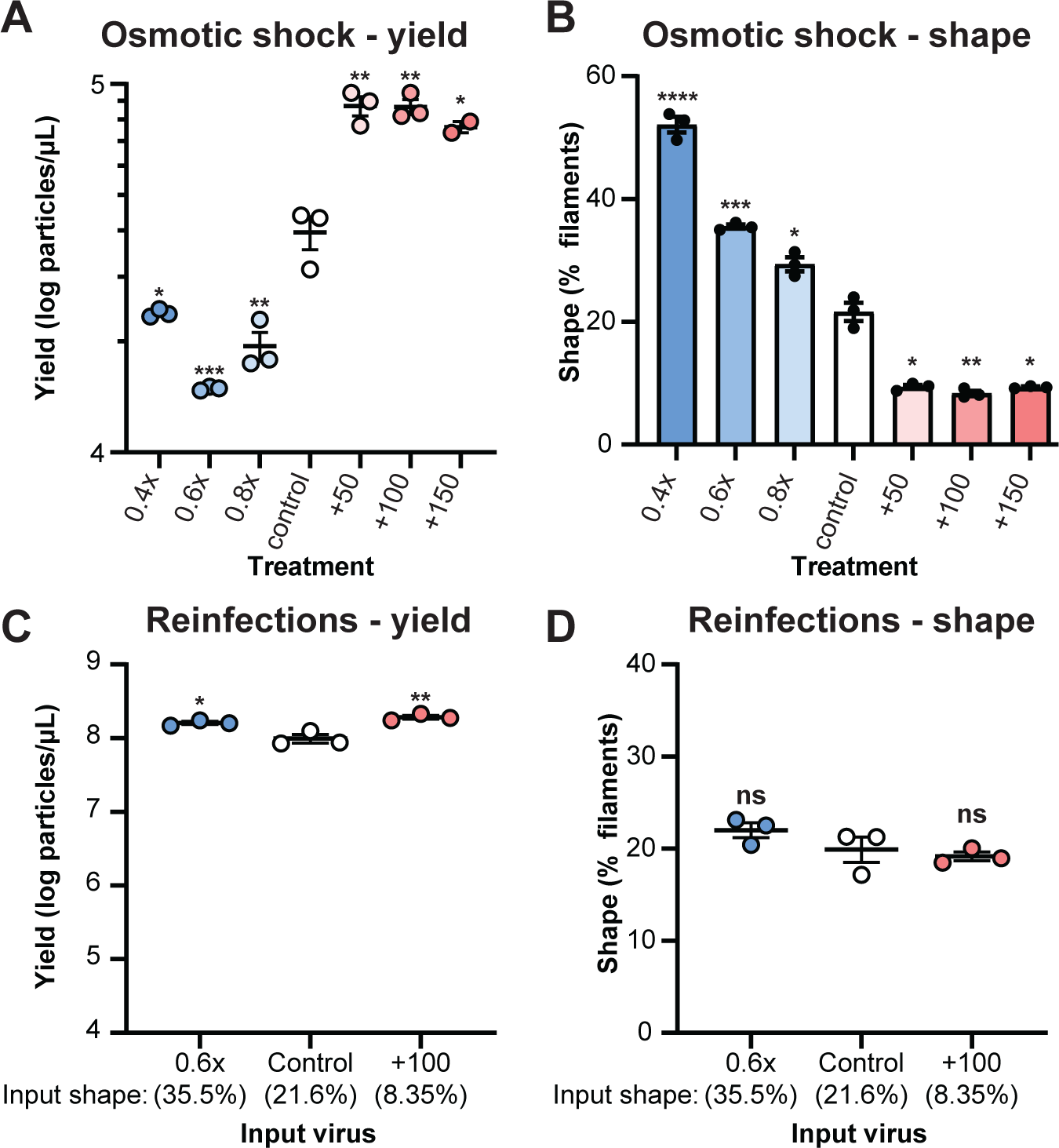
Osmotic Shock Alters Virion Assembly. MDCK cells infected with PR8 at MOI 0.3 for 20 hours were washed into media of varying osmotic strengths. **A**. Viral yields 30 minutes post media change. **B**. Shape of the samples in A. **C**. Yields 24 hours post infection of MDCK cells with the virus produced in A at 3 virions/cell. **D**. Shape of the samples in C. All plots show mean and S.E.M. *p<0.05, **p<0.01, ***p<0.001, ****p<0.0001 by unpaired t-test relative to control.

We aimed to specifically evaluate the effects of membrane tension on virion morphology in the context of infection. To do this, we performed single-cycle infections in the presence of IAV inhibitors targeting viral processes that occur either before or after endocytosis. Viral endocytosis is expected to increase membrane tension by depleting the cell membrane and increasing cell volume (Supplemental File 1). Inhibition with antibody Sb-H36-26 prevents attachment and therefore acts prior to endocytosis. The antibody MEDI8852 allows attachment and entry by endocytosis but prevents fusion, and baloxavir allows all entry steps but inhibits the viral polymerase (Figure 6A). We found that inhibitors that permit virion endocytosis (MEDI8852 and baloxavir) resulted in a greater proportion of filaments than inhibitors that act prior to endocytosis (Sb-H36-26, Figure 6B). This suggests that at least in the context of attenuated infection (i.e. low virion yields) differences in the extent of preceding endocytosis alter the shape of newly assembling virions.

**Figure 6.**
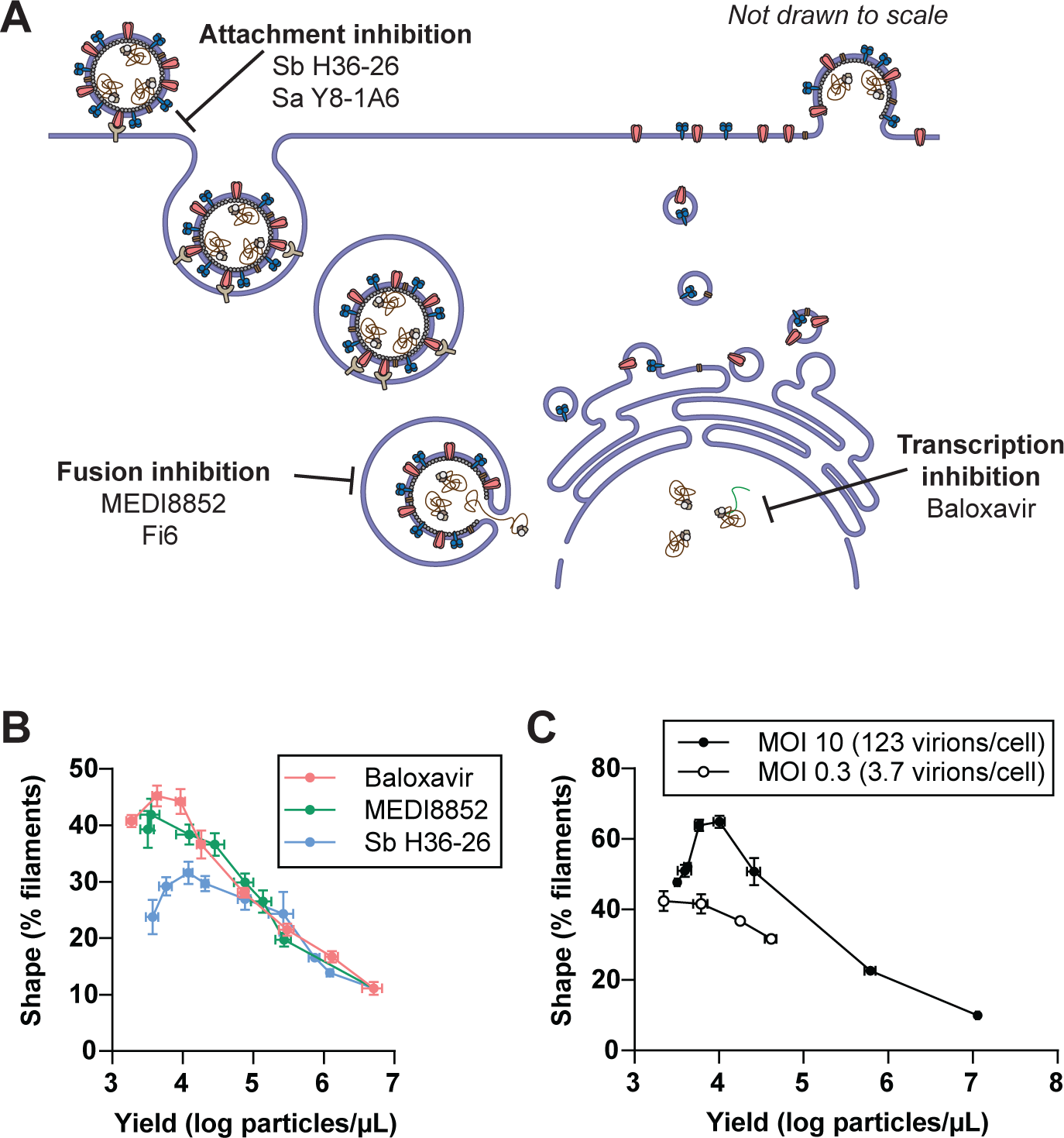
Filament Assembly Responds to Membrane Tension. **A.** Diagram outlining mechanisms of various inhibitors within the influenza viral life cycle. **B**. Shape vs. yield 24 hours post-infection of MDCK cells with PR8 at MOI 3 in the presence of a range of inhibitor concentrations (see also Figure S9). **C**. Shape vs. yield 24 hours post-infection of MDCK cells with PR8 at MOI 10 or MOI 0.3, with baloxavir included to normalize yields. All plots show mean and S.E.M.

To confirm that changes in cellular endocytosis at low yields affect viral assembly, we performed single-cycle infections at 123 and 3.7 input virions/cell (MOI 10 and MOI 0.3) while inhibiting the viral polymerase with baloxavir to assay a similar range of reduced yields. We thus expect much greater viral entry and endocytosis at high MOI, but similar levels of viral protein production, export, and virion formation. In these conditions, we found that high-MOI infections resulted in a greater proportion of filaments (Figure 6C), which is opposite of what we observed for uninhibited infections, and is consistent with a model that membrane tension induced by increased endocytosis can promote filament assembly at low virion outputs.

## Discussion

This work reveals that influenza A virion shape is dynamic, rather than a fixed property of a given strain. Assembly of filaments is responsive to the environment of the infection, including the identity of the host, the extent of infection, and the presence of antiviral antibodies. We also show that modulating host-membrane tension either artificially (by osmotic shock) or through normal viral processes (virion endocytosis) directly affects the shape of assembling virions. Our data, along with the advantage filaments have in animal infections, suggest that virion assembly may be a critical checkpoint during infection that allows influenza to sense and respond to environmental challenges.

Strong selection pressures could have driven evolution of pleomorphy in influenza. In otherwise unchallenging conditions, spherical particles are more efficient at entry and require fewer resources. Basal production of a small proportion of filaments would allow persistence once immune challenge becomes present. A virus that dynamically responds to this challenge by increasing the proportion of filaments would have an immediate advantage in subsequent rounds of infection. Enabling further infection cycles under immune pressure then creates opportunities for true antibody-escape mutants to arise (Figure 7).

**Figure 7.**
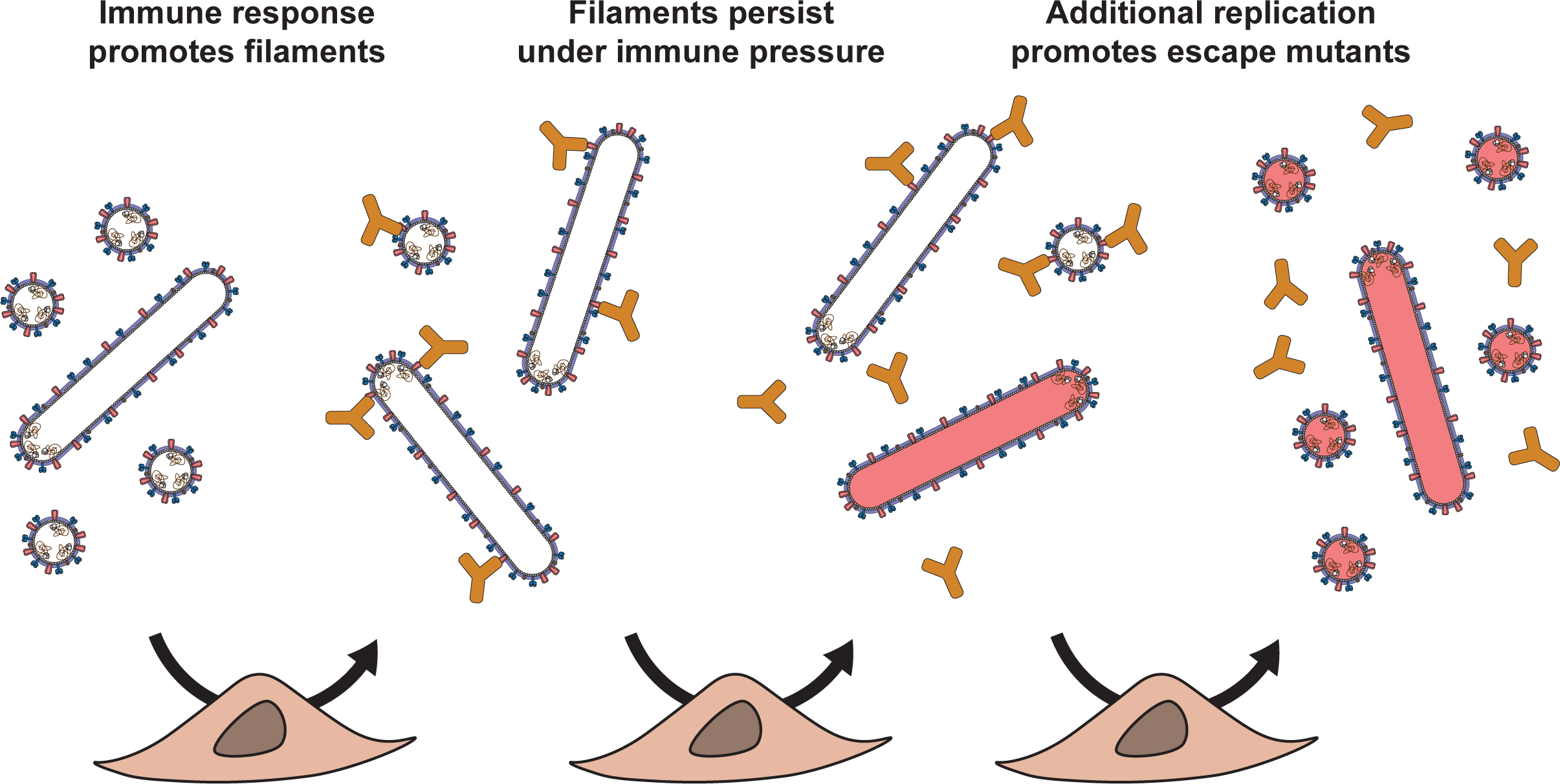
Filaments as a Mechanism for Viral Persistence. Model illustrating how dynamic virion shape distribution may promote viral persistence. A predominantly spherical population infects an animal cell, triggering an immune response. Antibody presence induces filament production. These filaments persist under immune pressure, allowing further replicative cycles, and possibly generating variants that evade antibody inhibition.

Dynamics in virus pleomorphy could enable adaptive mutations to arise when most needed, such as when a strain is introduced to a new host. For such spillover to occur, the virus must overcome two major barriers: receptor binding in the environment of the new host and polymerase adaptations^33^. When virus is first introduced to a new host, filaments would have higher likelihood of entering cells, despite unadapted receptor binding. We found that inhibiting the viral polymerase with boloxavir increases the proportion of filaments. Thus, during spillover, we would expect more filaments to be produced in the earliest replication cycles while the polymerase is unadapted. This would further increase the likelihood of the infection spreading in the non-cognate receptor environment. These key early replicative cycles under selection would then drive genetic adaptation to the new host.

We observe that general inhibition of infection increases the proportion of filaments - with effects observed from reduced MOI, inhibition of fusion, and inhibition of the viral polymerase. While the environment may be the source of such inhibition, in the case of antibody presence or transmission to a new host, it could also result from attenuation of the virus. Influenza A is a quickly mutating strain and can acquire spontaneous deleterious mutations. Furthermore, true resistance mutations to broadly neutralizing antibodies or inhibitors are expected to attenuate targeted functions^34,35^. The intrinsic ability of influenza to shift to filament production may dampen the negative effects of such a mutation. This would increase the chances for a secondary compensating mutation arise, and thus expand the possible evolutionary pathways to traverse valleys of low fitness.

Phenotypic adaptation to attenuating mutations could also be a confounding factor in studies of influenza morphology. An attenuating mutation might be misinterpreted as a mechanistic shape mutant if phenotypic tuning isn’t considered. Thus, proposed shape mutants must be carefully examined to ascertain whether they are truly reporting on the mechanisms of filament assembly.

Our osmolarity experiments implicate host-cell membrane tension in regulating virion shape outcomes. In uninfected cells, membrane tension is maintained in part through a balance between endocytosis and exocytosis^36^. Both of these processes are hijacked by viral processes, potentially affecting virion morphology. Virion endocytosis increases tension, export of viral transmembrane proteins in secretory vesicles decreases tension, and virion budding increases tension. The shape of produced virions in our time courses suggest that viral processes may be the dominant contributors to cell membrane tension changes. Internalization of 72 virions (approximately what is expected in our MDCK PR8 High MOI experiments) would decrease cell surface area by several percent (Supplemental File 1). Our time course infections initially produce filamentous virions, consistent with the expected tension present as a result of this endocytosis. As viral protein production and secretion increases, the shape of produced virions becomes more spherical, possibly in response to decreased tension. Finally, late in the time courses when budding may be the dominant process, filaments gradually increase (Figure 1, S3-4). In this way our shape observations parallel the expected membrane tension conditions in the cells over the time course of a high multiplicity infection.

It is initially counterintuitive that a high membrane tension would promote filament assembly, rather than simply reducing the assembly of smaller spherical particles. We propose that under high tension, the fission step of assembly is inhibited, but extension remains possible. This could potentially lead to additional elongation of assembling virions. When we subject infected cells to an osmotic shock that lowers membrane tension, we observe a rapid burst of particle release (Figure 5A). These particles are equivalently infectious as particles from untreated infections (Figure 5C) suggesting that they may have been fully assembled virion buds, stalled at some decision step between fission and extension. It is also possible that fission and extension are not always in competition, but that an elongation mechanism is engaged specifically when tension is high. This mechanism is unidentified, but one observation is that the matrix protein (M1) forms an ordered helical oligomer in filaments, a structure not seen in spherical virions. This model parallels the process of endocytosis; high membrane tension can stall pit formation, leading to the recruitment and oligomerization of actin or bar domains^37,38^. Interestingly, the involvement of these proteins can result in larger final vesicles compared to those formed under low tension. Perhaps like cells, viruses have evolved specific mechanisms to adapt to a high-tension environment.

Our MOI experiments (Figures 2, 6C) further link membrane trafficking to virion shape. We observed that the burst size is correlated to shape outcome. As MOI increases, the output/input ratio decreases, leading to a decrease in filament production. This aligns with our tension model if increased secretory pathway activity relieves tension more than increased tension from viral budding. Decreasing the output while holding the input constant (MEDI8852/Baloxavir, Figure 6B) results in more filament production, and decreasing the input while keeping the output constant (Figure 6C) leads to additional filaments production. These observations support the tension model.

It is unclear how changes in endocytosis during viral internalization translate to shape changes 24 hours later (Figure 6). While endocytosis is expected to increase membrane tension, it is surprising that these biophysical effects would persist over such time scales. An intriguing possibility could be that transient changes in tension induce a cellular response that then affects viral assembly. This would imply that influenza has even more sophisticated mechanisms of sensing and responding to host membrane biology. Sensing input particle numbers could benefit the virus; high input suggests that infections are productive and spheres are more efficient, while low inputs suggest infections are challenged and that filaments may yield advantages. Further modulation of shape during assembly would exaggerate effects further. The virus-infected cell can be viewed not merely as a passive site of viral replication, but as a living viral organism that leverages the complex capabilities of the cell to advantage the infection.

It is not clear by what mechanisms antibody-induced changes to shape could occur. Previous work has focused on genetic determinants of virion morphology. However, we find that while the viral genome contributes to virion-shape outcomes, it is not a strict determining factor as commonly assumed (compare Figures 1E and S3B), suggesting additional mechanisms must be at play. While general to all antibodies tested, it does not appear that effects are non-specific. Effects clearly vary based on the epitope, and also between cell types. Also, not every antibody that binds to viral proteins elicits assembly effects. For example, Ca H2-4B1-14 is highly neutralizing to PR8 in MDCK cells, suggesting it is capable of binding to HA, yet has no shape effects when present during assembly. In contrast, O19 binding to M2 has almost no ability to neutralize infection, yet does confer shape effects in assembly. Furthermore, there were no shape effects from a pan human MHC class I antibody in Calu3 cells (Figure S7). Our combined results suggest that the effects we see result from specific effects of antibody binding to cells, which we discuss in what follows.

Perhaps antibody effects are due to changes in membrane biophysics, akin to osmotic shock. The binding of antibodies to surface proteins could be affecting membrane tension or fluidity in a way that favors filament production. Perhaps antibody binding to or clustering viral proteins triggers their endocytosis, akin to growth factor endocytosis upon dimerization. While viral proteins aren’t normally expected to undergo endocytosis, we observe evidence that antibodies are depleted from infection supernatants. Specifically, when the antibody Cb H9-D3-4R2 was added to untreated virion supernatants, it induced severe clumping, yet the same concentration included in active infections did not. This could suggest that antibody-protein complexes are internalized, but further experiments are needed.

The environment of infection significantly impacts shape outcomes, as evidenced by the stark differences observed in virion shape produced from MDCK and Calu3 cells. A notable example is the spherical morphology of XUdorn when propagated in Calu3 cells. This finding challenges the prevailing belief that XUdorn is inherently filamentous, thereby emphasizing the role of host-virus interactions in determining virion shape. An additional characteristic of Calu3 cells is their insensitivity to changes in MOI, as these alterations do not significantly affect the shape of virions produced from them (Figure 1E). Further, the burst size changes only slightly with varying MOI, consistent with the proposed direct relationship between burst size and the shape of resulting virions. Yet another distinctive feature of Calu3 cells is the relatively lower per-particle infectivity of both PR8 and XUdorn compared to that in MDCK cells (Figure S2C). In summary, our data suggest that host-virus interactions play a substantial role in defining viral characteristics such as shape and infectivity, and that the environment of infection can induce different outcomes even for the same virus strain.

Flow virometry has enabled us to probe previously inaccessible aspects of viral biology. The technique resolves individual particles in a way that is not captured by the HA-assay and at a rate far exceeding that of electron microscopy. Moreover, flow virometry features ease of training, few limitations of sample concentration and purity, and minimal sample processing before performing measurements. While this paper addresses the morphology of influenza A, flow virometry is readily extendable to other viruses. It can be used to understand pleomorphy in general. These benefits allow us to approach questions that are not realistically answered by traditional means such as how phenotypic changes to morphology play a role in viral adaptation. Future applications of this technique may also define critical steps in pandemic emergence such as spillover, host adaptation, and evolution of vaccine escape variants.

## Acknowledgements

We would like to thank Jon Yewdell’s Lab at the National Institutes of Health and Steve Harrison’s lab at Harvard University for providing hybridomas, antibody constructs, and/or antibodies. We thank Jon Yewdell, Gunther Hollopeter, and Ivan Kosik for their comments to aid in preparing this manuscript. We acknowledge support from the NIH Director’s New Innovator Award 1DP2GM128204 and the NSF MRSEC DMR-1420382. This publication is based on research supported by The G. Harold and Leila Y. Mathers Charitable Foundation. Funding for this study was in part provided by the Divisions of Intramural Research of the National Institute of Allergy and Infectious Diseases. The content of this publication does not necessarily reflect the views or policies of the U.S. Government, nor does the mention of trade names, commercial products, or organizations imply endorsement by the U.S. Government.

## Author Contributions

Conceptualization, E.A.P. and T.I.; Methodology, E.A.P., S.D.P., and T.I.; Validation, E.A.P. and T.I.; Formal Analysis, E.A.P., A.J.W., S.D.P., and T.I.; Investigation, E.A.P., A.J.W., S.D.P., and T.I.; Resources, E.A.P., A.J.W., N.B., Z.L., and T.I.; Writing - Original Draft, E.A.P.; Writing - Review & Editing, E.A.P., A.J.W., S.D.P., and T.I.; Visualization, E.A.P., A.J.W., S.D.P., N.B., and T.I.; Supervision, E.A.P. and T.I.; Project Administration, E.A.P. and T.I.; Funding Acquisition, T.I.

## Methods

### Reagents

#### Cells

Madin-Darby canine kidney (MDCK)-Siat1 cells (Sigma-Aldrich, 05071502), MDCK.2 cells (ATCC strain CCL-34), and human embryonic kidney (HEK)293T cells (ATCC) were propagated in DMEM (Cytiva) supplemented with 10% FBS (Atlas Biologicals). Calu-3 cells (ATCC) were propagated in EMEM (Wisent) supplemented with 20% FBS. HEK293F cells, a gift from S. C. Harrison (Harvard Medical School), were propagated in FreeStyle 293 Expression Medium (Thermo Fisher Scientific) and were used for the expression of HC19 IgG, MEDI8852 IgG and 14C2 IgG.

#### Antibodies

Purified Fi6, CR9144, O19, and W6/32 antibodies, hybridomas producing anti-influenza virus HA antibodies Sb H36-26, Sa Y8-1A6-6, Ca H2-4B1-14, and Cb H9-D3-4R2, and hybridomas producing anti-influenza virus NA antibodies NA1-1c1 and NA2-10e10 were gifts from Jonathan Yewdell (NIH/NIAID/LVD/CBS). Hybridoma producing anti-influenza virus M2 monoclonal antibody 65 was a gift from Xavier Saelens (VIB-UGent Center for Medical Biotechnology). Hybridoma producing anti-influenza virus NP monoclonal antibody HB65 was obtained from ATCC (H16-L10-4R5, 58696953). The expression vectors (modified pVRC8400) for HC19 and MEDI8852 IgG heavy and light chains were a gift from S. C. Harrison (Harvard Medical School). The sequence for F045 was previously described^39^. For 14C2 IgG, the variable heavy and variable light chain sequences from a single-chain variable fragment^40^ were incorporated into a mouse IgG expression plasmid. Expressed antibodies were produced by transient transfection of HEK293F cells with PEI (0.4 μg heavy chain DNA, 0.6 μg light chain DNA, 1.5 μg PEI, per 1 x 10^6^ cells) and harvested from the cell culture supernatant 7 d after transfection. Antibodies from hybridomas, HC19 IgG, MEDI8852 IgG, and 14C2 IgG were purified using protein G resin (Cytiva). TriHSB.2 plasmid was a gift from David Baker (University of Washington). TriHSB.2 was expressed by autoinduction in bacteria as described^31^. F045 and TriHSB.2 constructs contain C-terminal 6xHis tags and were purified by passage over a HisTrapFF column (GE Healthcare), followed by size exclusion chromatography using a Superdex 200 increase 10/300 column (Cytiva). Antibodies were stored in PBS or PBS containing 10% glycerol.

#### Antibody labeling

Lyophilized DyLight550 NHS ester dye (Thermo Fisher Scientific) or JF646 NHS ester dye (Janelia) was resuspended in anhydrous DMSO (Sigma-Aldrich). The labeling reaction was performed with a 100:1 (HC19 or Sb H36-26, DyLight550) or 20:1 (HB65, JF646) molar ratio of dye to antibody in 100 mM sodium bicarbonate buffer for 1 hour at room temperature, followed by passage over a Macro SpinColumn packed with G25 packing material (Harvard Apparatus) in PBS. Labeled antibody concentrations were calculated using the measured absorbance at 280 and 557 nm.

### Viruses

A/Puerto Rico/8/1934 (PR8) Wild-Type influenza virus and A/Udorn/307/1972 containing the A/Aichi/1968 (X31) HA segment (XUdorn)^41^ were passaged in Calu3 cells at multiplicity of infection (MOI) 0.002 with 1 µg/mL TPCK-trypsin (Sigma-Aldrich) in OptiMEM (Thermo Fisher Scientific). The HA and NA segments were sequenced to ensure correctness in stock viruses. Viruses used here were passaged once from these stocks, and behavior was indistinguishable in our assays. Viruses were titered by standard hemagglutination (HA) and plaque assays. Hemagglutination assays were performed with chicken red blood cells and plaque assays on MDCK.2 cells.

#### Infectivity assays

For each cell line/virus strain combination, infectivity (virions/IU) was calculated from the average result of infections at two MOIs, all of which was performed in triplicate. Calu3 cells were infected with 0.4 and 2 PR8 virions per cell or 0.5 and 2.6 XUdorn virions per cell. MDCK cells were infected with 0.12 and 0.6 PR8 virions per cell or 0.15 and 0.8 XUdorn virions per cell (see single-cycle infections below). 24 hours post-infection (h.p.i), cells were fixed and permeabilized as described previously^22^ and then stained with JF646-labeled HB65, which recognizes NP. The number of infected cell singlets was determined by flow cytometry (NP deriving from input particles binding to cells at t0 was not detected under these conditions). The percent of cells infected was 5 or less, ensuring infected cells resulted from a single infectious unit.

#### Virus purification

XUdorn viruses were purified through a 20% sucrose cushion (SC), then either by passage over a 20-60% sucrose gradient^41^ or by sequential centrifugation as previously described^22^. Three fractions were collected from the sucrose gradient, namely ‘Band’, ‘Smear1’, and ‘Smear2’. Band is a distinct band, and Smear 1 and 2 correspond to virion samples deriving from diffuse-virus (‘smeared’) regions of the gradient of progressively higher sucrose density. From the sequential centrifugation, influenza virus pellet (Pellet1-6) fractions and supernatant (Sup1-4) fractions were collected.

### Electron microscopy

For electron microscopy, ‘Pellet4’, ‘Smear1’, and ‘Smear2’ were adjusted to 2 × 10^4^ HAU per mL and ‘SC’ and ‘Band’ were adjusted to 1 × 10^5^ HAU per mL. As a precaution, viruses were pretreated with 25 μM rimantadine (M2 inhibitor, Sigma-Aldrich) for 30 min at room temperature to prevent fragmentation of the filamentous particles^42^. Phosphotungstic acid (2%) staining was performed as previously described^41^. Images were taken using the Philips Morgagni v.3.0 transmission electron microscope (80 kV) using an AMT NanoSprint5 camera. Particle-size measurements were performed using custom MATLAB codes as previously described^41^.

### Time course experiments

MDCK-Siat1 or Calu-3 cells were grown in 24-well plates until they formed confluent monolayers. The cells were washed twice with HBSS (Thermo-Fisher Scientific) before attachment of PR8 or XUdorn virus in OptiMEM at MOI 0.006 or 6 for 1 hour at room temperature. After attachment, unattached virus was removed and the cells were washed twice with HBSS. 300 µL infection media, with or without trypsin and with or without ammonium chloride, was added. 4 h.p.i., ammonium chloride was added to some wells. 10 µL samples were taken every 4 hours for 48 h.p.i. Infected-cell supernatant aliquots were analyzed by flow virometry. Samples could be stored at −80°C until analysis.

### Single-cycle infections

#### General

Cells were grown in 24-well plates until they formed confluent monolayers. After washing twice in HBSS, virus was added in 35 µL OptiMEM and incubated at room temperature for 1 hour with frequent shaking. After attachment, cells were washed twice with HBSS and 300 µL OptiMEM was added. Trypsin was omitted to restrict infection to a single cycle. Infections were incubated at 34°C with 5% CO_2_ and 100% humidity. Infected-cell supernatant aliquots were analyzed by flow virometry. Samples could be stored at - 80°C until analysis.

#### MOI experiments

MDCK-Siat1 cells were infected with PR8 virus at a range of MOIs (see single-cycle infections above). Infected-cell supernatants were collected 24 h.p.i.

#### Antibody experiments

MDCK-Siat1 cells were used for PR8, and Calu-3 cells for PR8 and XUdorn. PR8 or XUdorn virus and antibody dilutions were made in OptiMEM. MEDI8852, Fi6, CR9114, HC19, monoclonalAb65, 14c2, and O19 antibodies were used for both viruses. Antibodies used only for PR8 were Sa Y8-1A6-6, Sb H36-26, Ca H2-4B1-15, Cb H9-D3-4R2, NA2-1c1, and NA2-10e10. XUdorn-specific antibodies were F045 and TriHSB.2. Cells were infected (see single-cycle infections above) at MOI 0.6. For PRE and ALL, a range of the indicated treatment was included during the room-temperature incubation and first four hours of infection. At 4 h.p.i., the media was removed and the cells were washed with HBSS. New infection media containing untreated OptiMEM for PRE or OptiMEM with treatment for POST and ALL were added to the cells. Infected-cell supernatants were collected 24 h.p.i.

#### Inhibitor experiment (Figure 6B)

MDCK-Siat1 cells were infected with PR8 virus at MOI 3 (see single-cycle infections above). When included, Sb H36-26 and MEDI8852 were included during the room-temperature incubation and first four hours of infection. At 4 h.p.i., the media was removed and the cells were washed with HBSS. When included, Baloxavir marboxil (Medchemexpress) was added to the indicated concentration from a 875 µM stock in cell culture DMSO (Sigma-Aldrich) at the end of the room-temperature incubation.

#### Inhibitor experiment (Figure 6C)

MDCK-Siat1 cells were infected with PR8 virus at MOI 0.3 or 10 (see single-cycle infections above) with ranges of Baloxavir treatments. When included, Baloxavir was added at the end of the room-temperature incubation. Infected-cell supernatants were collected 24 h.p.i.

#### Transient antibody treatment experiment

MDCK-Siat1 cells were infected with PR8 virus at MOI 0.6 (see single-cycle infections above). At 20 h.p.i., cells were washed into a range of concentrations of MEDI8852. Samples of the supernatant were collected at 1, 2 and 4 hours post treatment with MEDI8852 and analyzed by flow virometry.

#### Osmolarity experiments

MDCK-Siat1 cells were infected with PR8 virus at MOI 0.6 (see single-cycle infections above). At 20 h.p.i., cells were washed into either OptiMEM diluted with ultrapure H_2_O (0.4x, 0.6x, 0.8x), or OptiMEM supplemented with NaCl (50, 100, 150mM). Samples of the supernatant were collected at 30 minutes post treatment and analyzed by flow virometry.

#### Reinfection experiments

MDCK-Siat1 cells were infected with PR8 supernatants from the osmolarity or antibody treatment experiments (see single-cycle infections above). Infections were performed at 3 virions/cell in 35 µL. After 1 hr attachment, cells were washed into fresh OptiMEM. At 24 h.p.i., samples of the supernatant were collected and analyzed by flow virometry.

### Clumping experiments

An untreated supernatant from the antibody experiments (see above) were mixed with antibodies at the highest concentrations used in the antibody experiments. Mixtures were incubated at 34°C for one hour, and then analyzed by flow virometry.

### Flow virometry

DyLight550-labeled Sb H36-26 IgG and HC19 IgG stock solutions were diluted to 11.85 nM and 50 nM, respectively, in 0.2% BSA and HNE20 (20 mM HEPES NaOH pH 7.4, 150 mM NaCl, and 0.2 mM EDTA). Infected-cell supernatants were undiluted or diluted up to 1:30 in HNE20 and combined 1:1 with antibody dilution in BSA. Sb H36-26 IgG and HC19 IgG were used to label PR8 and XUd viruses, respectively. Binding reactions were incubated at room temperature for 1 hour, then diluted 1:250 in HNE20. Flow virometry was performed using the CytoFLEX S platform (Beckman Coulter). Laser powers were 70mW for violet and 50mW for yellow. Gain values were set to 300 for violet and 1000 for yellow. Samples were triggered on violet side-scatter area (1000-2000 A.U. threshold) and RFP (300-500 A.U. threshold) and acquired for 600 seconds or 25,000 particles in the virion gate. Thresholds were optimized for each instrument. Unlabeled, concentrated virus preparations were triggered on VSSC and acquired after 2 minutes until 500,000 particles in the virion gate (Figure 1B, Figure S1). Virus samples at variable dilutions (1:10 to 1:40,000) were mixed with FluoSphere (Invitrogen) beads (170nm, 505/515) at 1:300 and acquired by flow cytometry (Figure S1). All flow virometry samples were prepared in HNE20.

