## Supplemental File 1 for "Influenza A Virus Infections Sense Host Membrane Tension to Dynamically Tune Assembly"

**Supplemental file 1. Estimation of surface area and volume change from virion endocytosis**

1. Assume an MDCK cell is a cylinder with a radius of 15µm and a height of 7µm^43^.

Cylinder surface area = 2πrh + 2πr^2^

**Cell surface area** = 2*π*15*7 + 2*π*15^2^ = **2073µm^2^**

Cylinder volume = πr^2^h

**Cell volume** = π*15^2^*7 = **4948µm^3^**

2. Assume a 150nm endocytic vesicle is needed to surround a 100nm spherical virion and the glycoproteins^44^. 150nm diameter = 0.075µm radius.

Sphere surface area = 4πr^2^

**Sphere endocytic vesicle area** = 4*π*0.075^2^ = **0.07069µm^2^**

Sphere volume = 4/3 * πr^3^

**Sphere endocytic vesicle volume** = 4/3 * π * 0.075^3^ = **0.001767µm^3^**

3. Filaments enter by macropinocytosis. Assume a macropinosome is 0.2-5µm in diameter^45^.

   **Macropinosome area minimum** = 4*π*0.1^2^ = **0.1257µm^2^**

**Macropinosome area maximum** = 4*π*2.5^2^ = **78.54µm^2^

Macropinosome volume minimum** = 4/3 * π * 0.1^3^ = **0.004189µm^3^**

**Macropinosome volume maximum** = 4/3 * π * 2.5^3^ = **65.45µm^3^**

4. High MOI MDCK/PR8 experiments were performed at ~123 virions per cell. Input virus was 24% filamentous (see Figure 1D, time 0).

   Area internalized with spheres = 123 virions/cell * 0.76 (fraction spheres) * 0.07069µm^2^
   = 6.608µm^2^

   Minimum area internalized with filaments = 123 * 0.24 * 0.1257µm^2^ = 3.711µm^2^
   Maximum area internalized with filaments = 123 * 0.24 * 78.54µm^2^ = 2319µm^2^
   Geometric mean of minimum and maximum = sqrt(3.711*2319) = 92.77µm^2^

   Total area internalized = 6.608µm^2^ + 92.77µm^2^ = 99.38µm^2^
   **Fraction of cell membrane** = 99.38/2073 = **4.79%**

   Volume internalized with spheres = 123* 0.76 * 0.001767µm^3^ = 0.1652µm^3^

   Minimum volume internalized with filaments = 123 * 0.24 * 0.004189µm^3^ = 0.124µm^3^
   Maximum volume internalized with filaments = 123 * 0.24 * 65.45µm^3^ = 1932µm^3^
   Geometric mean of minimum and maximum = sqrt(0.124*1932) = 15.48µm^3^

   Total volume internalized = 0.1652µm^3^ + 15.48µm^3^ = 15.65µm^3^
   **Fraction of cell volume** = 15.65/4948 = **0.32%**
